# Electrical and G-protein Regulation of CaV2.2 (N-type) Channels

**DOI:** 10.1101/2024.06.29.600263

**Authors:** Michelle Nilsson, Kaiqian Wang, Teresa Mínguez-Viñas, Marina Angelini, Stina Berglund, Riccardo Olcese, Antonios Pantazis

**Affiliations:** Division of Cell and Neurobiology (CNB), Department of Biomedical and Clinical Sciences (BKV), Linköping University, Linköping 581 85, Sweden; Department of Anesthesiology and Perioperative Medicine, David Geffen School of Medicine, University of California, Los Angeles; Los Angeles, CA 90095, USA; Department of Physiology, David Geffen School of Medicine, University of California, Los Angeles; Los Angeles, CA 90095, USA; Wallenberg Center for Molecular Medicine, Linköping University, Linköping 581 85, Sweden

## Abstract

How G-proteins inhibit N-type, voltage-gated, calcium-selective channels (Ca_V_2.2) during presynaptic inhibition is a decades-old question. G-proteins Gβγ bind to intracellular Ca_V_2.2 regions, but the inhibition is voltage-dependent. Using the hybrid electrophysiological and optical approach voltage-clamp fluorometry, we show that Gβγ acts by selectively inhibiting a subset of the four different Ca_V_2.2 voltage-sensor domains (VSDs I-IV). During regular “willing” gating, VSDs I and IV activation resemble pore opening, VSD III activation is hyperpolarized, and VSD II appears unresponsive to depolarization. In the presence of Gβγ, Ca_V_2.2 gating is “reluctant”: pore opening and VSD-I activation are strongly and proportionally inhibited, VSD IV is modestly inhibited while VSD III is not. We propose that Gβγ inhibition of VSD-I and -IV underlies reluctant Ca_V_2.2 gating and subsequent presynaptic inhibition.

## Introduction

N-type voltage-gated calcium channels Ca_V_2.2 are found at the presynaptic terminal of neurons in the central and peripheral nervous systems, where their activation initiates calcium-dependent neurotransmitter release (*1*) (Fig.1A). Ca_V_2.2 channels are renowned for their abundance in nociceptors and role in development and treatment of chronic pain (*1*). Neurotransmitters and neuromodulators such as noradrenalin, serotonin, GABA (*2*) and opioids (*3*) can inhibit Ca_V_2.2. This was initially demonstrated in 1978 in dorsal root ganglion (DRG) neurons (*2*) and was eventually ascribed to G-protein induced inhibition of Ca_V_2.2 (*1, 4*). The Gβγ complex directly inhibits Ca_V_2.2 opening, reducing calcium influx and subsequent neurotransmitter release (*1*) (Fig.1B).

**Fig. 1:**
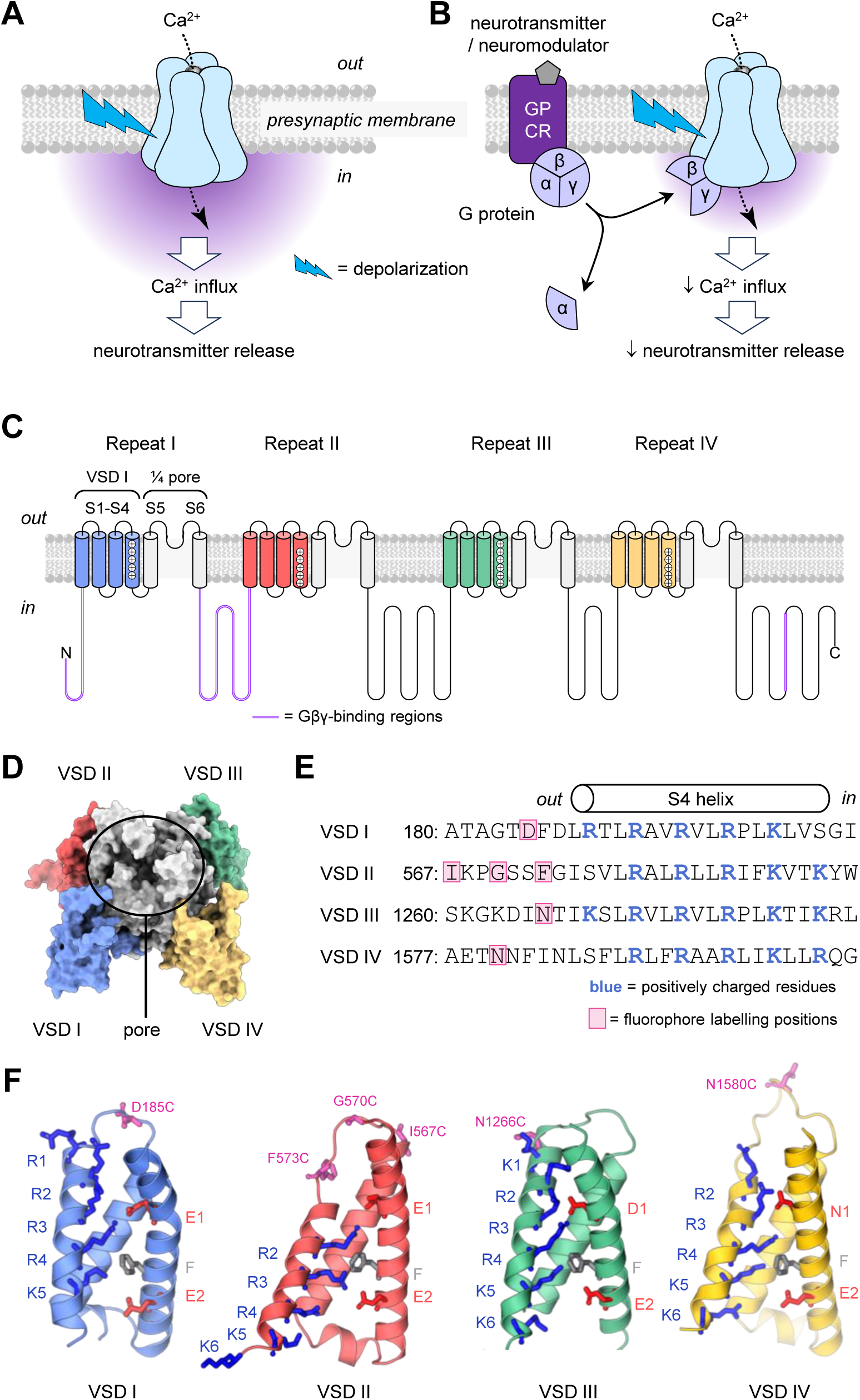
Ca_V_2.2 function, G-protein inhibition and structure. (**A**) Arrival of an action potential to the presynaptic terminus triggers activation of Ca_V_2.2 channels, Ca^2+^ influx and neurotransmitter release (*1*). (**B**) Neurotransmitters and neuromodulators can activate their cognate, presynaptic G-protein-coupled receptors (GPCRs) triggering G-protein hydrolysis and release. The Gβγ complex directly inhibits the voltage-dependent opening of Ca_V_2.2, reducing Ca^2+^ influx and neurotransmitter release (*1*). (**C**) Membrane topology of the CaV2.2 pore-forming subunit (α_1B_). It comprises four homologous repeats, each spanning the membrane six times. The first four transmembrane segments (S1-S4) assemble into a distinct voltage-sensor domain (VSD), while S5 and S6 from all repeats form the central pore. Gβγ is thought to bind to cytosolic loops regions (magenta): the N-terminus, the repeat-I-II loop and the C-terminus. (**D**) Structure of a human Ca_V_2.2-channel (*6*). (**E**) Amino-acid sequence diversity among the four Ca_V_2.2 repeats. Positively-charged residues (blue) drive VSD activation upon membrane depolarization. Pink boxes show cysteine-substitution positions for fluorescence labelling and voltage-clamp fluorometry. (**F**) Ribbon structures of VSDs I-IV (*6*). Pink residues show cysteine-substituted positions used for site-directed fluorescent labelling. Cobalt-blue residues are positively charged arginines (R) and lysines (K). Grey residues are phenylalanines (F) indicating the charge-transfer center. Red residues are negatively-charged counter-charges (CC) in S2. Note the differences in gating charges R1-K6 and that VSD II was resolved in a down-state (resting, gating charges below F).

The Ca_V_2.2 α_1B_ pore-forming subunit consists of four interlinked repeats (I-IV) that form a central calcium-conducting pore domain (transmembrane segments S5-S6 from each repeat) and four voltage-sensor domains (VSDs, segments S1-S4 from each repeat) that surround and control opening of the pore (*5–7*) (Fig.1C,D). The four VSDs differ in amino-acid composition and likely respond differentially to depolarization (*8, 9*). Binding sites for Gβγ have been identified in the N-terminus, Repeat I-II loop and C-terminus of Ca_V_2.2 (*10–12*) (Fig.1C). Curiously, G-protein inhibition is voltage-dependent. As described by Bean, G-protein inhibition changes calcium channel gating from *willing* to *reluctant*, shifting voltage-dependent activation towards depolarized potentials (*4*). G-proteins can also inhibit Ca_V_2.2 gating currents (*13–15*). Collectively, this points to an important role of the VSDs. Here, we present voltage-clamp fluorometry data illuminating activation of individual voltage-sensors and their inhibition by G-proteins—fundamental events controlling calcium-mediated transmitter release.

## Materials and Methods

### Molecular biology

The following constructs were used: human CACNA1B (hα1B; variant +e10a, +18a, Δ19a, +e31a, +e37b and +e46; Addgene #62574, a gift from Diane Lipscombe (*16*)), rabbit CACNA2D1 (α_2_δ-1; UniProt accession no. P13806), rat CACNB2a (β_2a_; UniProt accession no. Q8VGC3), rabbit CACNB3 (β_3_; UniProt accession no. P54286), human GNB1 (Gβ1; #140987, a gift from Bryan Roth (*17*)), human GNG2 (Gγ2; Addgene #67018, a gift from Catherine Berlot) and zebrafish voltage-sensitive phosphatase (DrVSP; a gift by Yashushi Okamura (*18*)). Genes in cell-expression vectors were subcloned to the Z-vector for *in-vitro* transcription and oocyte expression (*19*). Mutagenesis was performed using PfuUltra II Fusion High-fidelity DNA Polymerase (Agilent) or Q5 Site-Directed Mutagenesis Kit (New England Biolabs). To improve PCR yield with the GCrich hα1B template, reactions were supplemented as needed with 1-5% DMSO (Invitrogen) and/or 1M betaine (Thermo Scientific). Plasmid sequences were fully confirmed by sequencing. Finally, cRNA was transcribed *in vitro* using mMESSAGE mMACHINE T7 (ThermoFisher Scientific) or HiScribe T7 ARCA mRNA Kit (New England Biolabs), evaluated spectrophotometrically and by gel electrophoresis, and stored at −80°C.

### Oocyte preparation

Oocyte lobes were surgically removed from *Xenopus laevis* as approved by the Linköping Animal Care and Use Committee (Permit #1941), in accordance with national and international guidelines. Lobes were separated into clusters of 1-5 oocytes and the follicular layer was removed i) enzymatically with Liberase (Roche) and ii) mechanically. Both steps were done using an orbital shaker at 88 rpm at room temperature in Ca^2+^-free OR-2 solution (in mM: 82.5 NaCl, 2.5 KCl, 1 MgCl_2_, 5 HEPES, pH=7.0), first with Liberase for approximately 20 min, then in OR-2 for approximately 45-75 min. As needed, grade I defolliculated oocytes purchased from Ecocyte Bioscience were also used. Cells were stored in SOS (100 NaCl, 2 KCl, 1.8 CaCl_2_, 1 MgCl, 5 HEPES, pH=7.0) at 17°C.

Stage V-VI oocytes were injected (UMP3T-1, World Precision Instruments) at the equator with 50 nl cRNA mix containing the Ca_V_2.2 complex hα1B, α_2_δ-1, β_2a_ without or with Gβ1 and Gγ2. In PIP_2_ depletion experiments, hα1B, α_2_δ-1, β_3_ and DrVSP were co-injected. β_3_ was used as it augments Ca_V_2.2 sensitivity to PIP_2_ depletion (*20*). All subunits were injected at 0.3µg/µl. Oocytes were incubated for 3-4 days at 17°C in incubation solution (in final concentration: 50 % L-15 (Corning cellgro Leibovitz’s L-15), 47.5% H_2_O, 10% heat-inactivated horse serum (Gibco), 10^5^ u/l and 100 mg/l penicillin/streptomycin (Capricorn Scientific GmbH), 100 mg/l amikacin (Fisher BioReagents)). Of note, Gβγ modulation typically developed at day 4, even though Ca_V_2.2 channels readily expressed by day 3.

### Labelling

Before VCF recordings, oocytes were labelled for 7 minutes on ice with 20 μM thiol-reactive fluorophore MTS-TAMRA (MTS-5(6)-carboxytetramethylrhodamine, mixed isomers, Biotium) in a depolarizing solution (in mM: 120 K-methanesulfonate (MES), 2 Ca(MES)_2_, and 10 HEPES, pH=7.0). As described, in some VSD-II experiments, oocytes were labelled with 10 µM 6-TAMRA C6 maleimide (tetramethylrhodamine-6-maleimide C6, AAT Bioquest) or 100 μM Alexa-488 C5 maleimide (Thermo Fisher Scientific) on ice for 30 min. All stocks were 100 mM in DMSO.

### Electrophysiology

Voltage-clamp fluorometry (VCF) was performed in room temperature under cut-open oocyte Vaseline gap (COVG) as previously described (*21–23*). We used a CA-1B amplifier (Dagan Corporation). All signals were sampled with a Digidata 1550B1 digitizer and pClamp 11.2.1 software (Molecular Devices). The optical set-up comprised a BX51WI upright microscope (Olympus) with filters (all Semrock BrightLine: exciter: FF01-531/40-25; dichroic: FF562-Di02-25x36; emitter: FF01-593/40-25). The excitation light source was the M530L3 LED (530 nm, 170 mW, Thorlabs) driven by a Cyclops LED driver (Open Ephys). Fluorescence emission was acquired with a LUMP-LANFL 40XW water immersion objective (Olympus; numerical aperture = 0.8, working distance = 3.3 mm) and focused on a SM05PD3A Si photodiode (Thorlabs). Photocurrent was amplified with a DLPCA-200 current amplifier (FEMTO). For Alexa-488 experiments, the following filter-set was used (all Semrock BrightLine): exciter: FF01-482/35-25; dichroic: FF506-DI03-25X36; emitter: FF01-524/24-25. The light source was a Thorlabs blue LED (490 nm, 205 mW, M490L4). Current and fluorescence were sampled at 25 kHz and low-pass–filtered at 5 kHz.

Before mounting to the COVG chamber, each oocyte was injected with 100 nl 100 mM BAPTA•4K (Invitrogen), 10 mM HEPES, pH = 7.0, to prevent calcium-dependent Ca_V_2.2 regulation and oocyte currents. External solution (mM): 120 NaMES, 2 Ba(MES)_2_, 10 HEPES (pH=7.0). Internal solution (mM): 120 K-Glutamate, 10 HEPES (pH=7.0). Intracellular micropipette solution (mM): 2700 NaMES, 10 NaCl and 10 HEPES (pH=7.0).

Cells were held at −80 mV and stepped to a range of voltages for 50 ms when characterizing VSDs, or 20 ms when investigating G-protein modulation. In experiments investigating G-protein modulation, a pre-pulse facilitation protocol was used to confirm Gβγ modulation. (*24, 25*) Cells were held at −100 mV and the level of facilitation was determined at a 0 mV test pulse before and after a conditioning 100 ms pre-pulse at 100 mV. In cells co-injected with Gβγ, only cells displaying ≥50% facilitation were included in data analysis.

Action potential waveform was derived from a model based on recordings in rat DRG neurons at room temperature (*26*) in Matlab R2022a (MathWorks) using the *ode15s* differential equation solver. In action potential clamp experiments cells were held at −60 mV, as this was the cell resting potential.

### Analysis

Analysis was performed in Clampfit 11.2 (Molecular devices), Excel (Microsoft) and Matlab (MathWorks).

The Ca_V_2.2 tail current *I*_tail_ was obtained from the peak tail current, fitted to a Boltzmann equation

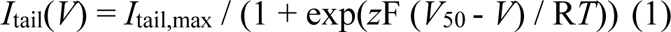

Where *V* was the membrane potential, *I*_tail,max_ was maximal *I*_tail_, *z* was the valence, *V*_50_ is half-activation potential and *R* and *F* the gas and Faraday constants, respectively.

Δ*F* values were obtained from the steady-state activation by the end of each pulse and fit to a Boltzmann equation

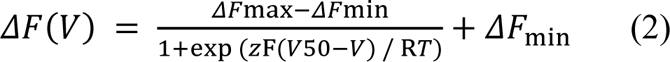

where Δ*F*max was the maximum fluorescence deflection and Δ*F*min the minimum fluorescence deflection.

Gβγ pre-pulse facilitation “Ppf” was measured as the ratio of the current after and current before a facilitating pre-pulse, at a 0-mV test pulse at 22 ms, when facilitated current approach a plateau. This protocol was performed in all Gβγ experiments, and only cells displaying at least 50% facilitation were further analysed.

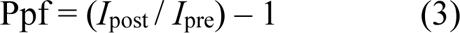

In Gβγ facilitation recordings of Δ*F*, facilitation was determined as above. The relative facilitation was calculated from the facilitation of the current in the same cell

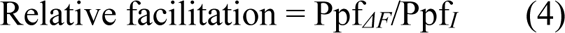

We generated a DRG-neuron action potential using the Choi and Waxman formulation (*26*), and used it as voltage command in our VCF set-up.

The conductance *G* was calculated as

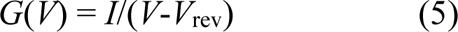

where *I* is the current across the trace, *V* the membrane potential and *V*_rev_ the reversal potential as determined from an *I*_tail_-*V* 50-ms pulse protocol in the same cell, obtained at the same *V*_rest_ (−60mV). *G*(*V*) was then normalized as

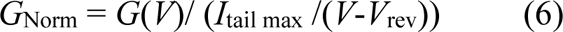

where *I*_tail max_ was the maximum tail current derived from the *I*_tail_-*V* protocol and *V* was the membrane potential (−60 mV).

The fluorescence was normalized by

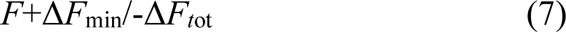

where *F* is the fluorescence across the trace, Δ*F*_min_ is the minimum fluorescence as determined from an Δ*F*-*V* 50 ms protocol in the same cell, obtained at the same *V*_rest_ (−60 mV), and Δ*F*_tot_ is the total change in fluorescence determined by the Δ*F*-*V* protocol. In fluorescence traces from VSD IV where the fluorescence deflection is positive, −Δ*F*_min_ was used. The normalized *G* or Δ*F* at *V*_rest_ was determined at −60 mV and the normalized *G* or Δ*F* peak was determined as the maximum conductance or Δ*F*.

### Statistics

All tests were two-tailed unpaired Student’s *t*-tests, except for the statistics on the relationship between VSD and conductance inhibition in action-potential-clamp experiments (Fig 5C): in this case, the *p* value calculated using two-sample Kolmogorov-Smirnov tests.

### Protein Structures

Ca_V_2.2 channel structures with Protein Data Bank accession number: 7MIY (*6*) were rendered using The Protein Imager (*27*). Movie S1 was made on Blender 3.3.0 (The Blender Foundation).

## Results

### Ca_V_2.2-VSDs have diverse responses to depolarization

To determine the voltage-sensing properties of the Ca_V_2.2 VSDs, we implemented voltage-clamp fluorometry (VCF) (*21, 28, 29*). VCF enables the optical dissection of the individual VSDs in conducting Ca_V_2.2 macromolecular complexes (α_1B_/α_2_δ-1/β_2a_) under physiologically relevant conditions. Ca_V_2.2 channel complexes were expressed in *Xenopus laevis* oocytes. MTS-TAMRA, an environment-sensitive fluorophore was conjugated to a cysteine substituted at the S3-S4 linker of each VSD (Fig.1E), reporting VSD activation as fluorescence deflections (Δ*F*). The membrane potential was controlled with the cut-open-oocyte Vaseline gap (COVG) method (*22, 30, 31*).

Ca_V_2.2 cysteine variants of VSDs I (D185C), III (N1266C), and IV (N1580C) generated reliable Δ*F* upon voltage-dependent activation (Fig.2A), while maintaining wild-type-like pore-opening voltage dependence (Fig.2B). Δ*F* from VSDs I and III was quenched upon depolarization (downward deflections), whilst Δ*F* from VSD IV was unquenched (upward deflection) (Fig.2A). All VSDs responded differently to voltage (Fig.2), as suggested by their distinct amino-acid composition and structural poses (Fig.1E,F) (*6, 7*). VSD II did not generate any Δ*F*, comparable to channels without substituted cysteines, regardless of the positions tested (Figs.2A & S1). This suggests that VSD II does not respond to depolarization. This is a remarkable finding that explains why recent structures of Ca_V_2.2, acquired in the absence of an electric field (0 mV), revealed S4_II_ in a resting conformation (*6, 7*) (Fig.1F).

**Fig. 2:**
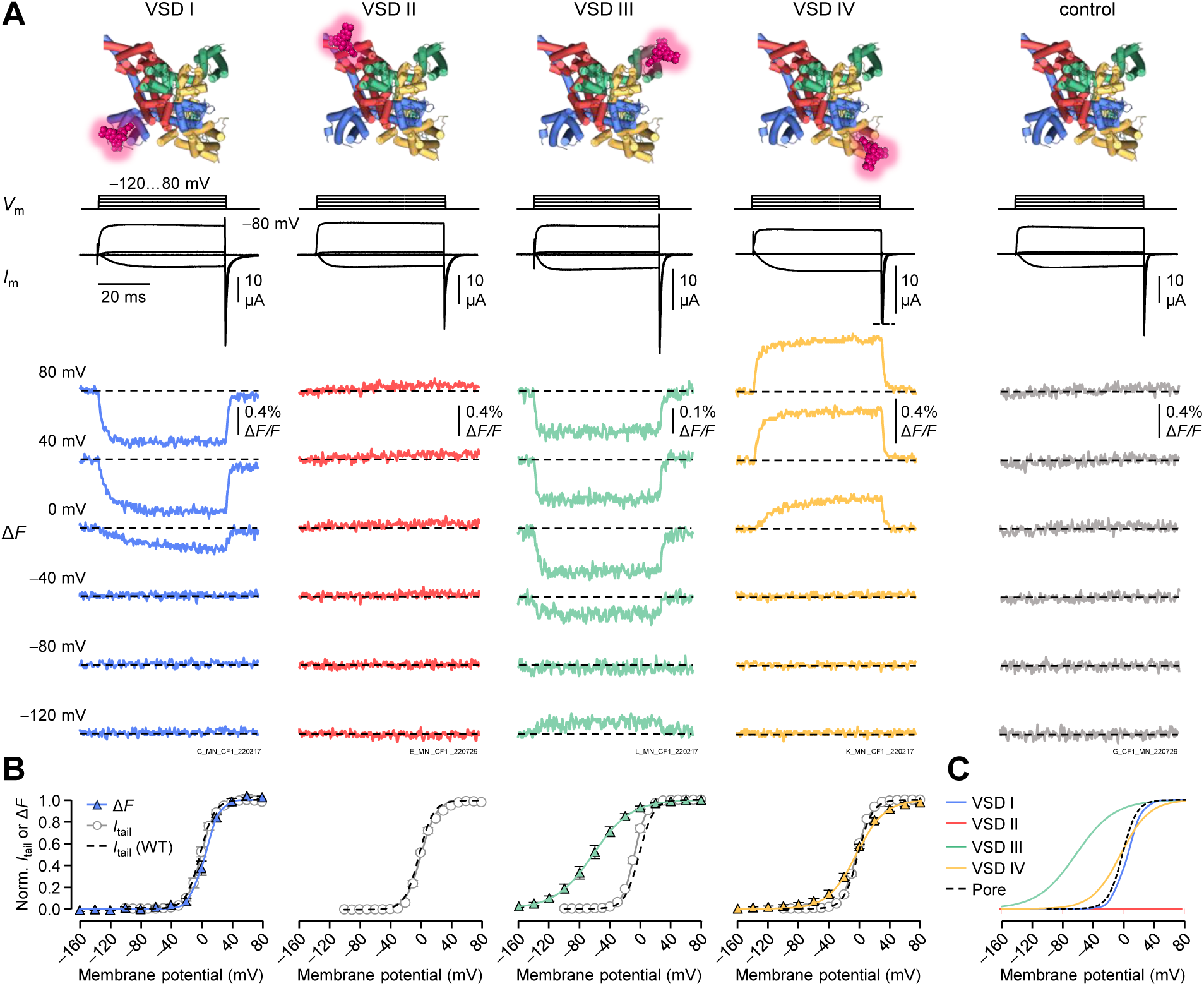
Distinct voltage-sensor activation properties of the Ca_V_2.2 VSDs. (**A**) Voltage-clamp fluorometry recordings in Ca_V_2.2. Membrane potential (*V*_m_) steps, corresponding current (*I*_m_) traces and simultaneously acquired fluorescence deflections (Δ*F*) from each VSD, as well as control channels without substituted Cys. Note that VSD II in this and other experimental conditions (Figs.S1,S2) showed no evidence of voltage-dependent activation. (**B**) Normalized tail-current-voltage (*I*_tail_-*V*) and Δ*F*-voltage (Δ*F*-*V*) curves showing the voltage-dependent pore opening and VSD activation, respectively. The grey dotted line shows the wild-type (control) *I*_tail_-*V* curve for comparison (*V*_50_ = 1.0±1.7 mV, *z* = 2.9±0.2 *e*_0_, *n*=6). Boltzmann distribution parameters: VSD I *V*_50_ = 4.7±2.6 mV, *z* = 3.0±0.3 *e*_0_, *n*=5. VSD III *V*_50_ = −62.8±5.6 mV, 1.1±0.1 *e*_0_, *n*=7. VSD IV *V*_50_ = −5.9±1.5 mV, *z* = 1.6±0.2 *e*_0_ *e*_0_, *n*=12. Error bars represent SEM. (**C)** The voltage-dependence curves from (b) shown together for comparison, and to accentuate the functional diversity of the four Ca_V_2.2 VSDs. As VSD is apparently voltage-insensitive, its voltage-dependence is represented by a flat red line.

VSD-I activation and pore opening appeared to be coupled, as observed by the closely overlapping fluorescence deflection-voltage (Δ*F*-*V*, blue) and tail-current-voltage (*I*_tail_-*V,* black) curves (Fig.2B,C). Specifically, VSD I activated with a half-activation potential *V*_50_ = 4.7±2.6 mV and had an apparent voltage sensitivity *z* = 3.0±0.33 *e*_0_ (*n*=5). This was comparable to pore opening of this construct (*V*_50_ = 0.091±2.6 mV, *z* = 2.8±0.19 *e*_0_, *n*=5).

VSD II did not respond to voltage changes. We tested three, uniformly spaced, labelling positions in the S3_II_-S4_II_ linker: I567C, G570C and F573C (Fig.1E). Despite high functional expression, no Δ*F* could be observed, even at extreme voltages (−160 to 160 mV; Figs.2A & S1). To prevent any occurrence of stable quenching, we removed a nearby tryptophan (W564F), as in a previous VCF study on BK channels (*32*), but this made no difference (Fig.S1). We also tested labelling with fluorophores with a longer stalk, 6-TAMRA-C6-maleimide (*33*) or Alexa-488 C5-maleimide, but this did not generate any Δ*F* (Fig.S1). Initial structures of Ca_V_2.2 hinted that S4_II_ might be stabilized in the down state by a PIP_2_ molecule (*6, 7*) and PIP_2_ is known to modulate Ca_V_2.2 (*34*). Thus, we depleted PIP_2_ using the voltage-sensitive phosphatase DrVSP, as previously described (*20*), but still did not observe Δ*F* (Fig.S2). Subsequent structures of the Ca_V_2.3 channel also showed S4_II_ in a down state, in the absence of PIP_2_ (*35, 36*), suggesting that the “locked down” S4_II_ is a feature of the Ca_V_2 family independently of PIP_2_.

VSD III activated at strikingly negative potentials and had a half-activation potential of −62.8±5.6 mV, and low apparent voltage sensitivity at 1.1±0.1 *e*_0_ (*n*=7) (Fig.2B,C). As such, *ca.* 50% of Ca_V_2.2-VSD III would be active at resting membrane potentials. At −20 mV, 4% of channels were open, while 90% of VSD III were activated.

Finally, VSD IV had a half-activation potential of −5.9±1.5 mV, close to that of pore opening, and a low apparent voltage sensitivity of 1.6±0.2 *e*_0_ (*n*=12) (Fig 2B,C).

### VSD I responds more strongly than VSDs III and IV to pre-pulse facilitation

With the additional dimension of individual voltage-sensor resolution now available from VCF, we explored the mechanistic details of G-protein inhibition of Ca_V_2.2. To recapitulate Ca_V_2.2 inhibition by G-proteins in our experimental paradigm, we co-expressed the Ca_V_2.2 complex (α_1B_/α_2_δ1/β_2a_) together with the human Gβ1 and Gγ2 subunits. A hallmark of Gβγ inhibition is “pre-pulse facilitation” (*24, 25*). This describes the current increase observed following a strongly depolarizing pre-pulse during Gβγ inhibition, and is thought to be mediated by the transient unbinding of Gβγ and relief of Gβγ inhibition (*25*). When we optically tracked Ca_V_2.2 VSD activation in the presence of Gβγ, we found that the VSDs facilitated to a different extent (Fig.3): VSD I had the most prominent facilitation, VSD III did not respond to pre-pulse facilitation and VSD IV had an intermediate response. This revealed that GPCR signalling indeed modulates the Ca_V_2.2 VSDs, in a VSD-selective manner.

**Fig. 3:**
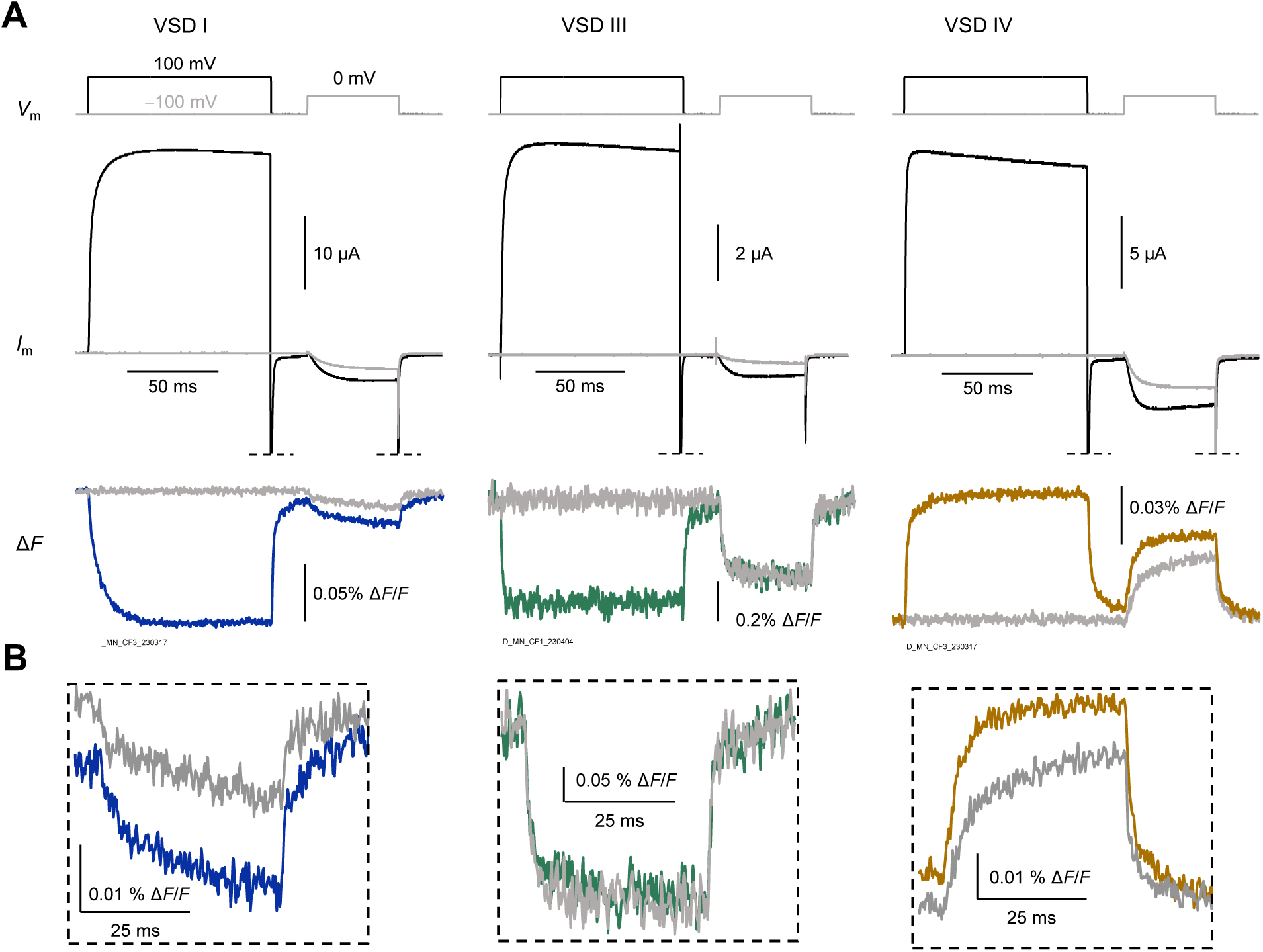
Distinct voltage-sensor facilitates upon pre-pulse facilitation in the presence of Gβγ. (**A**) Exemplary recordings of VSD pre-pulse facilitation in cells expressing α_1B_/α_2_δ-1/β_2a_/Gβ1/Gγ2. Top: Steps of membrane potential (*V*_m_) i) without pre-pulse (grey) and ii) with 100 ms facilitating pre-pulse at 100 mV (black), both followed by a test pulse at 0 mV. Middle: corresponding facilitating barium current (*I*_Ba_). Bottom: fluorescence deflections (Δ*F*) from VSD I, III and IV without (grey) and with (blue, green and yellow, respectively) facilitating prepulse. (**B**) Close-up of VSDs I, III and IV Δ*F* during the 0-mV test pulse.

### Gβγ makes VSDs I and IV “reluctant” to activate

To further determine Gβγ-inhibition of the VSDs, we studied their voltage-dependent activation in the absence or presence of Gβγ. To prevent relief of Gβγ inhibition (dependent on time and voltage), we used shorter (20 ms) activation pulses. As expected, Gβγ shifted the *V*_50_ of Ca_V_2.2 pore opening towards positive potentials, Δ*V*_50_= 14±0.8mV, *p* = 3e−7 (Fig.4A,B).

**Fig. 4:**
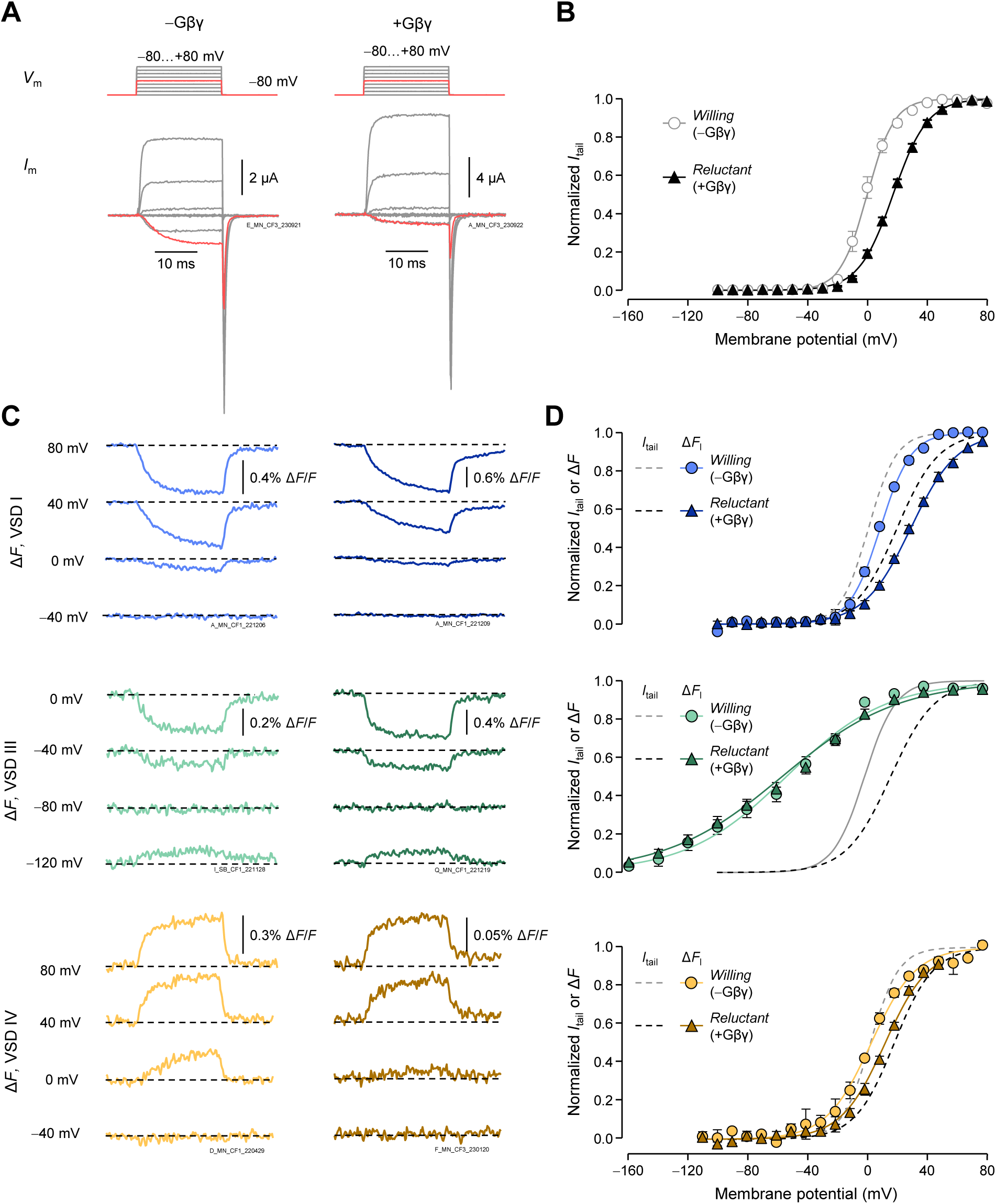
G-protein-induced inhibition of VSDs I and IV. (**A**) Steps of membrane potential (*V*_m_) and corresponding current traces without (−Gβγ, “willing”) or with Gβ1 and Gγ2 (+Gβγ, “reluctant”) from channels labelled in VSD-I. (**B**) Normalized (*I*_tail_-*V*) curves showing the voltage-dependent opening of wild-type Ca_V_2.2: −Gβγ: *V*_50_ = 3.3±0.9 mV, *z* = 3.2±0.2, *n* = 9; +Gβγ: *V*_50_ = 17±0.8 mV, *z* = 2.3±0.1 *e*_0_, *n* = 7. (**C**) Representative fluorescence deflections (Δ*F*) from VSDs I, III and IV in the presence or absence of Gβγ. (**D**) Normalized Δ*F* - voltage (Δ*F*-*V*) curves showing voltage-dependent activation of VSD I, III and IV. VSD I −Gβγ: *V*_50_ = 8±2 mV, *z* = 2.8±0.5 *e*_0_, *n* = 5; +Gβγ: *V*_50_ = 28±2 mV, *z*=1.8±0.2 *e*_0_, *n* = 7. VSD III −Gβγ: *V*_50_ = −54±7 mV, *z* = 0.8±0.1 *e*_0_, *n* = 6; +Gβγ: *V*_50_ = −54±5 mV, *z* = 0.7±0.03 *e*_0_, *n* =9. VSD IV −Gβγ: *V*_50_ = 1.5±1.6 mV, *z* = 2.1±0.3 *e*_0_, *n* = 10; +Gβγ: *V*_50_ = 14±1 mV, *z*=1.9 ± 0.2 *e*_0_, *n* = 10. Black solid line (−Gβγ) and black dotted line (+Gβγ) show *I*_tail_-*V* curves from the corresponding cysteine variant that Δ*F* was collected from. Error bars are SEM.

We discovered that VSD-I activation is strongly inhibited by Gβγ. Gβγ shifted the VSD-I *V*_50_ by 21±1.0 mV, *p* = 8e−8, similar to channel opening in the same cells (Fig.4C,D). To evaluate whether the effects on VSD-I activation was mediated *via* the canonical Gβγ binding sites, we introduced point-mutation R54A, previously shown to abolish G-protein inhibition (*37*). Indeed, Gβγ inhibition of both the pore opening and the VSD-I activation was eliminated (Fig.S3). In contrast to VSD I, VSD III was not modulated by Gβγ, Δ*V*_50_ = −0.1±5.3 mV, *p* = 0.99 (Fig.4C,D). VSD IV displayed modest inhibition by Gβγ compared to VSD I, Δ*V*_50_= 13±1 mV, *p =* 1E−6 (Fig.4C,D).

### VSD-I activation and pore opening are strongly and proportionately inhibited by Gβγ during DRG action potentials

Ca_V_2.2 is the predominant synaptic calcium channel in nociceptors (*38–41*) and is important in pain sensitivity and development of chronic pain (*42–44*). To investigate VSD operation under a physiological stimulus, we implemented action-potential clamp with a waveform characteristic of small, unmyelinated nociceptive DRG neurons (*26, 41, 45*). At the resting membrane potential (*V*_rest_, −60 mV) channels were closed (Fig.5A,B). In the case of VSD-I-labeled channels, the peak macroscopic conductance was observed during the plateau phase of the action potential, reaching 17±1% of the maximum, as normalized by a standard “*I*_tail_-*V*” protocol on the same cell. In the presence of Gβγ, the peak conductance was significantly reduced by approximately half, to 8±2% (*p* = 0.004; Fig.5C).

**Fig. 5:**
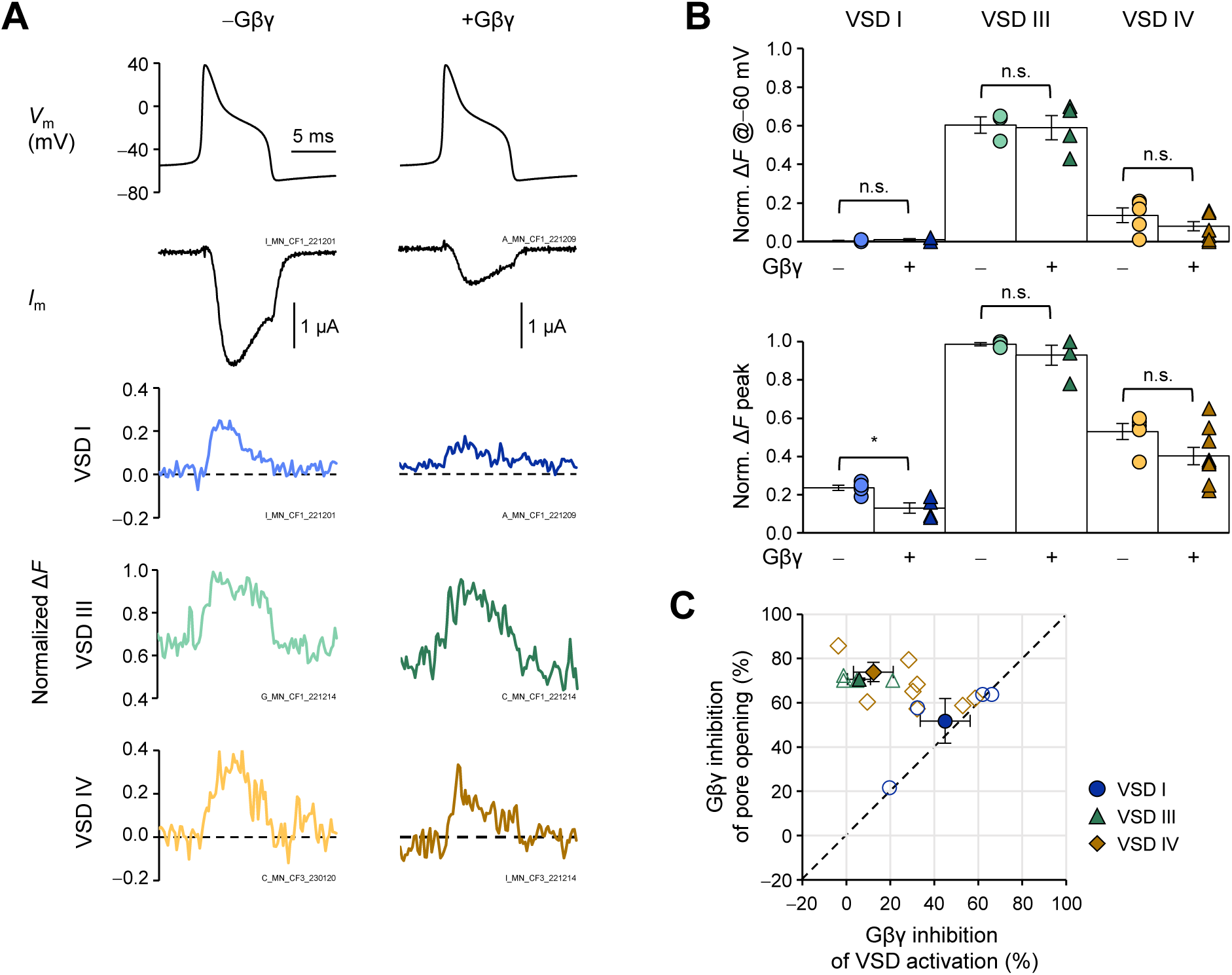
VSD I-activation and pore opening are strongly and proportionally inhibited by Gβγ during a DRG action potential. (**A**) Top: Action-potential clamp (*V*_m_) in cells expressing α_1B_/α_2_δ-1/β_2a_ without (−Gβγ “willing”) or with Gβ1 and Gγ2 (+Gβγ, “reluctant”). Middle: corresponding current traces. Bottom: normalized fluorescence deflections (Δ*F*) from VSDs I, III and IV. (**B**) Normalized Δ*F* calculated from (a), and a Δ*F*-*V* curve obtained from the same cell. Top: Normalized Δ*F* at resting membrane potential (*V*_rest_, −60 mV). VSD I −Gβγ: 0.0±0.0 %, *n* = 5; +Gβγ: 0.0±0.0 %, *n* = 4. VSD III: −Gβγ: 60±4 %, *n* = 3; +Gβγ: 59±6 %, *n* = 4, *p* = 0.88. VSD IV: −Gβγ: 14±4 %, *n* = 5; +Gβγ: 8±2 %, *n* = 9, *p* = 0.21. Bottom: Normalized peak Δ*F* VSD I: −Gβγ: 24±1 %, *n* = 5; +Gβγ: 13±3 %, *n* = 4, *p* = 0.007. VSD III: −Gβγ: 99±1 %, *n* = 3; +Gβγ: 93±5 %, *n* = 4, *p* = 0.4. VSD IV: −Gβγ: 53±4 %, *n* = 5; +Gβγ: 40±5 %, *n* = 9, *p* = 0.09. Tests were two-tailed, unpaired *t*-tests. n. s. = non-significant. * = significant, *p* ≤ 0.05. (**C**) Gβγ inhibition of pore opening (conductance) plotted against Gβγ inhibition of VSD activation. Open symbols represent individual experiments, filled symbols are means. The dashed line represents 1:1 inhibition between pore opening and the voltage sensors. VSD-I inhibition correlated well with conductance inhibition (*p* = 1.0). On the other hand, inhibition of VSDs III and IV did not correlate with G inhibition (*p* = 0.011 and 4.9E−4, respectively). Tests were two-sample Kolmogorov-Smirnov type. Error bars are SEM. VSD = voltage-sensor domain.

At *V*_rest_, VSD I was not activated, whilst already 60±4% of VSD III and 14±4% of VSD IV were active (Fig.5A,B). These fractions did not significantly change in the presence of Gβγ (VSD I: *p* = 0.33; VSD III: *p* = 0.88; VSD IV: *p* = 0.21).

Peak VSD activation was defined as the maximum amplitude of the Δ*F*, normalized by a standard “Δ*F*-*V*” protocol. Similar to pore opening, VSD I reached 24±1% activation and was significantly reduced to 13±3% in the presence of Gβγ (*p* = 0.007, Fig.5). VSD III reached maximal activation independently of Gβγ (−Gβγ; 98±10%; +Gβγ: 94±5%;*p* = 0.4). Finally, 53±4% of VSD IV activated at peak, which was not significantly different in the presence of Gβγ (40±5%, *p* = 0.09). To better illustrate the timing of VSD activations and pore opening, the activity traces were used to annotate the Ca_V_2.2 structure in Movie S1.

## Discussion

Our most recent knowledge on the voltage-sensor domains of Ca_V_2.2 came from the atomic structures resolved by cryogenic electron microscopy, effectively at 0 mV (*6, 7*). However, the dynamic responses of these structures to electrical signals and their regulation by G-proteins were unknown. Our optical investigation of the Ca_V_2.2 VSDs under physiologically relevant conditions revealed that these domains exhibit distinct voltage-sensing properties and regulation by G-proteins. In summary, we found that: i) VSDs I, III and IV activate with distinct voltage-dependencies, likely contributing in different ways to channel opening; ii) in our tested conditions, VSD II does not respond to changes in the membrane potential; iii) Gβγ sets VSDs I and IV in a “reluctant” mode of activation.

### The Ca_V_2.2 voltage-sensing apparatus

VSD I activation and pore opening are exhibit very similar voltage dependence (Figs.2,4,5), suggesting that this domain is strongly coupled to pore opening. In favour of this interpretation, action-potential clamp protocols revealed that VSD I activation and pore opening are coupled in both time and voltage (Fig.5, Movie S1). Both achieved maximal activation- or open-probability of roughly 20% during an action potential. Indeed, introducing a negative charge in Ca_V_2.2 S3_I_ , which might inhibit VSD-I activation, shifted *V*_50_ of pore opening by over +10 mV (*46*).

VSD II may not be pertinent for channel opening, as discussed below. VSDs III and IV both activate within physiologically relevant potentials. VSD III in particular appears to be tuned to sensing the resting membrane potential. Voltage-clamp recordings and stimulation with an action-potential waveform showed that 60% of VSD III are already active at resting membrane potential, when channels are closed (Figs.2,4,5). This anticipates that Ca_V_2.2 channels have at least two closed states: one with VSD III in the resting conformation, and another with VSD III in the active conformation. This disimplicates VSD-III activation from directly driving pore opening. VSD IV has a voltage-dependence approaching that of pore opening, although the latter saturates at more negative potentials than VSD IV activation (Fig.2). One interpretation is that VSDs I, III and IV are all required for opening of the pore, where VSD III activation is the first VSD to transition to an active state and VSDs I and/or IV are limiting factors for pore opening.

We did not observe any fluorescence deflections from VSD II despite testing multiple positions and conditions, even at extreme levels of depolarization, hyperpolarization and overexpression (Fig.2, Figs.S1,S2). Our findings, together with recent structural evidence, support a role of VSD II alternative to voltage sensing. In 2021, two independent studies both revealed Ca_V_2.2 channels with S4_II_ in a down state in the absence of electric field (0 mV), while S4_I_, S4_III_ and S4_IV_ were in an up-state (*6, 7*). Later structures of the closely related Ca_V_2.3 channel also demonstrated resting VSD II states (*35, 36*). Moreover, Ca_V_2.3 channels can undergo voltage-dependent opening upon neutralization of S4_II_ gating charges, with only a modest shift in *V*_50_ (*36*). VCF has revealed that VSD II does undergo voltage-dependent activation in Ca_V_1-channels (*8, 9*) and accordingly, S4_II_ in resolved structures of Ca_V_1 and Ca_V_3 channels is in an active conformation (*47–49*). We propose that a voltage-insensitive VSD II is a signature of Ca_V_2-family channels, where it may serve as a static structural element of the channel. Perhaps under different regulatory regimes or in other splice variants, VSD II can undergo voltage-dependent activation and contribute to Ca_V_2.2 opening and regulation.

### GPCR tuning of Ca_V_2.2 voltage-dependence

We found that VSD I is strongly inhibited by Gβγ, in proportion to pore-opening inhibition, whilst VSD IV is modestly affected and VSD III is not (Figs.4,5). Accordingly, VSD I activation is also facilitated most strongly by pre-pulse facilitation, while VSD IV shows more modest facilitation and III is unaffected (Fig.3). This hierarchy of VSD modulation by Gβγ suggests that Gβγ directly interacts with VSD I and potentially VSD IV, to prevent activation and subsequent opening. In single-channel recordings, Gβγ inhibition manifests as the emergence of low-open-probability (*P*_O_), “reluctant” openings, and delayed high-*P*_O_ openings (*50*). Consolidating this information and our results, we propose that Gβγ acts by inhibiting VSD-I-driven channel-opening (delaying high- *P*_O_ events), allowing the emergence of inefficient openings by VSD IV (low-*P*_O_ events). Indeed, the previously known Gβγ binding sites may position Gβγ in close proximity to VSD I (Fig.1C) (*10–12*). Gβγ-mediated stabilization of the VSD-I resting conformation may be due to electrostatic interactions. Indeed, introducing a negative charge in S3_I_ (G177E) to inhibit VSD-I activation resulted in similar effects as G-protein inhibition: Ca_V_2.2 opening becomes intrinsically reluctant, and sensitive to pre-pulse facilitation (*46*). Neutralizing the R2 gating charge of S4_I_ in the closely related Ca_V_2.1-channel also reduced G-protein inhibition (*51*). An interaction with VSDs I and IV in a down state could account for the preferential closed-state G-protein inhibition of Ca_V_2.2 (*52, 53*).

Ca_V_2.2 is an attractive target for pain management as it has been implicated both in the development and treatment of chronic pain (*1, 42, 44*). Indeed, the inhibition of VSDs I and IV poses an interesting way of regulating Ca_V_2.2 channels. For example, a drug that mimics this natural form of inhibition—a *Gβγ mimetic*—could be a promising next-generation analgesic, specifically targeted to inhibit VSDs I or IV. Indeed, since these domains are transmembrane, such compounds could be developed to act from the extracellular side. Tuning ion-channel activation by modulating VSDs have been shown both for toxins, drug-like compounds, and endogenous molecules (*54*).

In conclusion, we show that Ca_V_2.2-VSDs respond differently to voltage and that this asymmetry extends to their modulation by other proteins. Gβγ, the result of GPCR signalling in the presynaptic terminal, specifically acts by preventing the activations of VSDs I and IV.

## Acknowledgments

We thank members of the Pantazis group for useful discussions and members of the Elinder, Liin and Pantazis groups for oocyte preparation.

## Funding

Lions Forskningsfond Ph.D. support (M.N.)

NIH/NIGMS R35GM131896 (R.O.)

start-up funds from the Linköping University Wallenberg Center for Molecular Medicine / the Knut and Alice Wallenberg Foundation (A.P.)

Hjärnfonden (The Swedish Brain Foundation) grants FO2022-0003 and FO2023-0025 (A.P.)

Vetenskapsrådet (The Swedish Research Council) grants 2019-00988 and 2022-00574 (A.P.)

## Author contributions

Conceptualization: M.N., R.O., A.P

Methodology: M.N., T.M.-V., A.P

Investigation: M.N., K.W., T.M.-V., M.A., S.B

Visualization: M.N., A.P

Funding acquisition: M.N., R.O., A.P

Project administration: M.N., A.P

Supervision: M.N., A.P

Writing – original draft: M.N.

Writing – review & editing: All

---

**Fig. S1:**
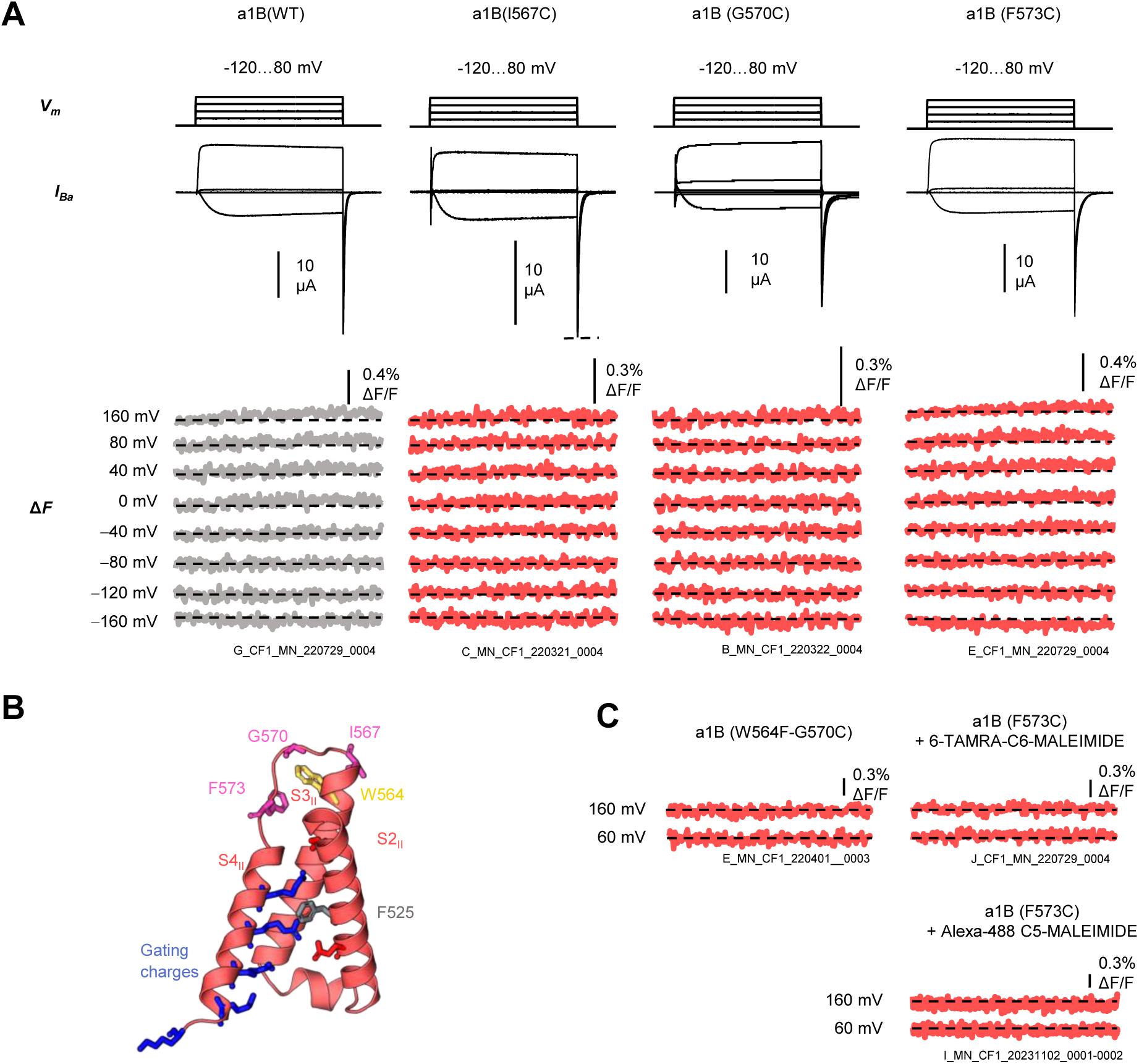
VSD II labelling conditions. **(A)** Top: voltage-step protocol, mid: exemplary current responses (*I*_Ba_), bottom: exemplary fluorescence deflections (Δ*F*) after staining with MTS-TAMRA. **(B)** Structural representation of Ca_V_2.2-VSD II (*6*), cysteine labelling positions highlighted in pink (I567C, G570C, F573C) and removed tryptophan in yellow (W564F). **(C)** Extended labelling conditions: removal of a potentially quenching tryptophan (W564F), labelling with 6- TAMRA-C6-MALEIMIDE or Alexa-488 C5-MALEIMIDE

**Fig. S2:**
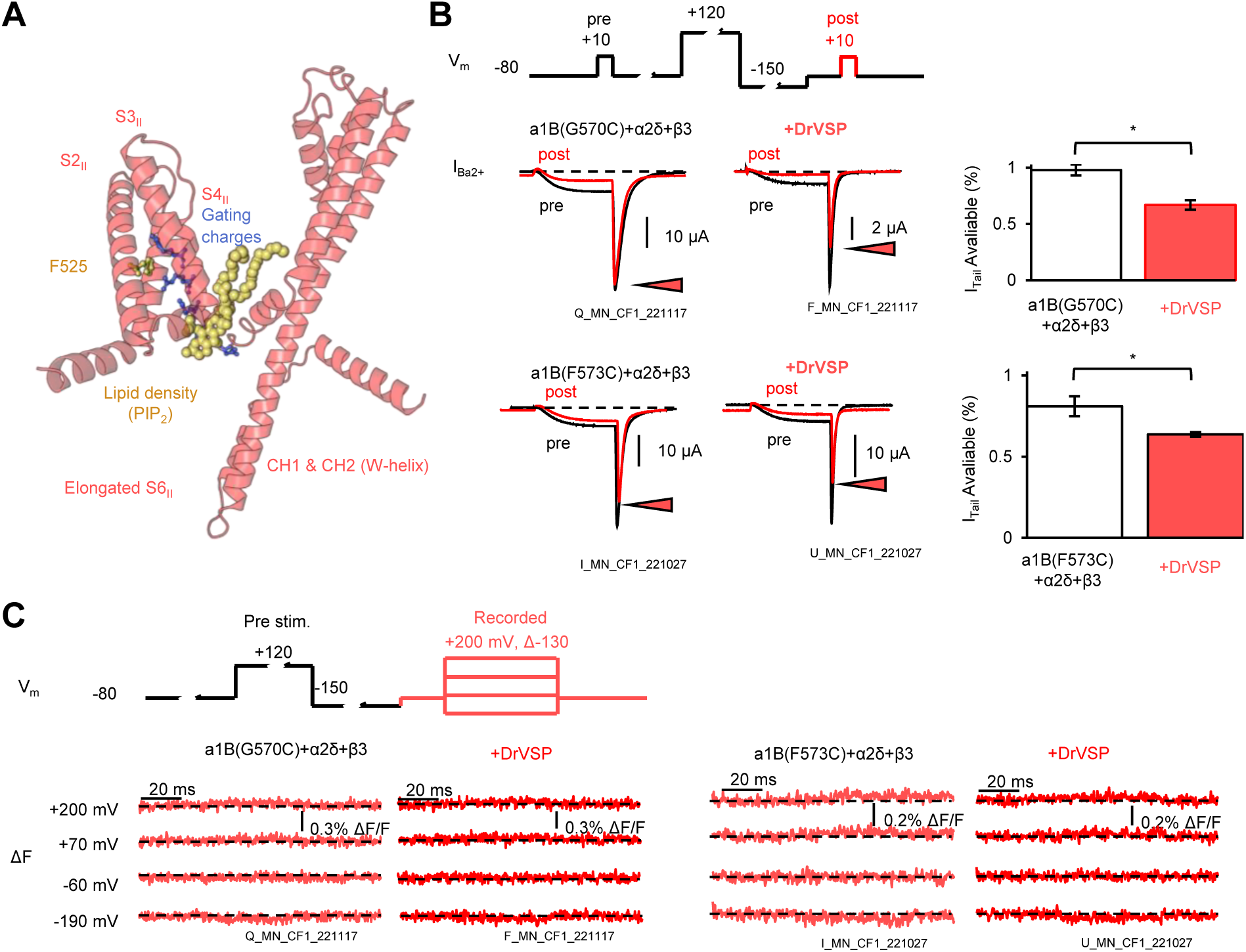
DrVSP-induced PIP_2_ depletion to relieve Ca_V_2.2-VSD II. **(A)** Structural representation of Ca_V_2.2 repeat II showing S4 in a down-state, potentially stabilized by a lipid density (PIP_2_) (*6*). **(B)** Top: Depletion protocol. Pre-test pulse at 10 mV, 10 ms, pre-stimulating pulse to 120 mV, 1000 ms, inactivation removal pulse to −150 mV, 400 ms, and post-test pulse at 10 mV. Bottom: avaliable tail current before (pre) and after (post) a PIP_2_-depleting pulse in abscence or presence of DrVSP. **(C)** Top: Depletion protocol followed by 50 ms test-pulses to various voltages, as indicated. Bottom: examplary fluorescence deflections (Δ*F*) in abscence or presence of DrVSP. Bars are mean ± SEM, * indicates *p*<0.05.

**Fig. S3:**
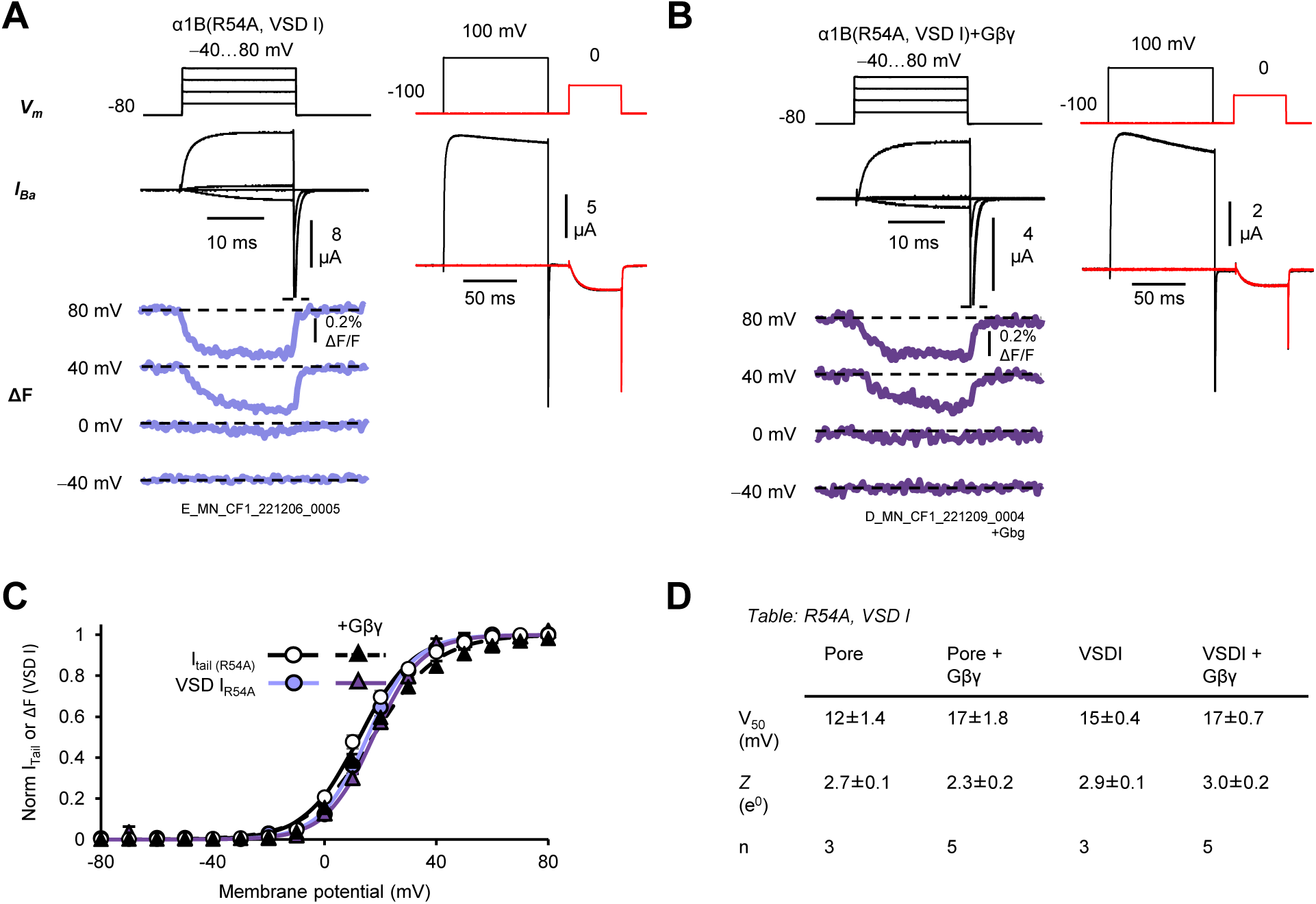
R54A prevents Gβγ inhibition of Ca_V_2.2-VSD I. **(A)** Left: voltage-dependent activation of VSD I, right: pre-pulse facilitation **(B)** as in A, but in the presence of Gβγ. **(C)** Voltage-dependent channel opening (I_tail_) and fluorescence deflection (Δ*F*) from VSD I. **(D)** Summary of voltage-dependent properties.

**Legend for Movie S1:** Structural dynamics of Ca_V_2.2 during activation by a DRG action potential. Distinct functional domains of the human Ca_V_2.2 channel structure (*6*) were rendered so that their activities are shown as changes in brightness. The data were collected using VCF with action-potential clamp, using the DRG action-potential waveform, in the absence of G-protein regulation (Fig.5A, left panels). Pore opening (conductance) is shown in purple, while the activations of VSDs I, III and IV is shown in blue, green and orange, respectively. VSD II is shown as having constant 0 brightness (no activation), as expected from its apparent lack of voltage sensitivity (Figs.2, S1, S2).

